# Patterns of MHC-dependent sexual selection in a free-living population of sheep

**DOI:** 10.1101/2020.11.18.387332

**Authors:** Wei Huang, Jill G. Pilkington, Josephine M. Pemberton

## Abstract

The MHC is one of the most polymorphic gene clusters in vertebrates and play an essential role in adaptive immunity. Apart from pathogen-mediated selection, sexual selection can also contribute to the maintenance of MHC diversity. MHC-dependent sexual selection could occur via several mechanisms but at present there is no consensus as to which of these mechanisms are involved and their importance. Previous studies have often suffered from limited genetic and behavioural data and small sample size, and were rarely able to examine all the mechanisms together, determine whether signatures of MHC-based non-random mating are independent of genomic effects or differentiate whether MHC-dependent sexual selection takes place at the pre- or post-copulatory stage. In this study, we use Monte Carlo simulation to investigate evidence for non-random MHC-dependent mating patterns by all three mechanisms in a free-living population of Soay sheep. Using 1710 sheep diplotyped at the MHC class IIa region and genome-wide SNPs, together with field observations of consorts, we found sexual selection against a particular haplotype in males at the pre-copulatory stage and sexual selection against female MHC heterozygosity during the rut. We also found MHC-dependent disassortative mating at the post-copulatory stage, along with strong evidence of inbreeding avoidance at both stages. However, results from generalized linear mixed models suggest that the pattern of MHC-dependent disassortative mating could be a by-product of inbreeding avoidance. Our results therefore suggest that while multiple apparent mechanisms of non-random mating with respect to the MHC may occur, some of them have alternative explanations.

## Introduction

Major histocompatibility complex (MHC) genes encode cell surface proteins that present pathogen-derived peptide to T cells to activate the adaptive immune response and are one of the most variable loci across the vertebrate genome. Although pathogen-mediated balancing selection is believed to be the major force shaping MHC diversity, sexual selection could also contribute to the maintenance of MHC diversity (Jordan and Bruford 1998, Penn 2002, Milinski 2006, Radwan, Babik et al. 2020). A potential role for MHC-dependent sexual selection has been assessed theoretically (Ejsmond, Radwan et al. 2014) and documented in empirical studies (Winternitz, Minchey et al. 2013, Kamiya, O’Dwyer et al. 2014, Winternitz, Abbate et al. 2017). MHC-dependent sexual section could act through both intrasexual selection via male-male competition and through intersexual selection including mate choice based on the partner’s MHC constitution (additive benefits) or based on the MHC compatibility (non-additive benefits).

More specifically, MHC-dependent sexual selection could occur via three non-mutually exclusive mechanisms: selection could favour particular MHC alleles or haplotypes, MHC diversity or MHC compatibility. First, individuals with specific MHC alleles could be favoured by intrasexual or intersexual selection and such a pattern has been reported in several studies (Ekblom, Saether et al. 2004, Eizaguirre, Yeates et al. 2009). Second, some studies have reported that individuals with higher diversity or heterozygosity at the MHC were favoured as partners (Landry, Garant et al. 2001, Richardson, Komdeur et al. 2005, Cutrera, Fanjul et al. 2012, Winternitz, Promerova et al. 2015). Third, in terms of MHC compatibility, MHC-dependent disassortative mating has been reported, which could maximise the MHC diversity of offspring (Schwensow, Eberle et al. 2008, Setchell, Charpentier et al. 2010, Hoover, Alcaide et al. 2018, Han, Sun et al. 2019). Several studies have also demonstrated that MHC genes could be a cue for kin recognition enabling avoidance of mating with relatives through MHC-associated odours (Grob, Knapp et al. 1998, Penn 2002, Milinski 2006). However, as excessive expression of MHC molecules could lead to depletion of the mature T-cell repertoire and elevated risk of autoimmune disease (Migalska, Sebastian et al. 2019), it has also been suggested that individuals may optimise rather than maximise MHC diversity (Reusch, Haberli et al. 2001, Griggio, Biard et al. 2011, Rekdal, Anmarkrud et al. 2019). Fourth, some studies have found mate choice favouring MHC-similar mates and explained such patterns as local adaption to endemic pathogens (Bonneaud, Chastel et al. 2006, Bichet, Penn et al. 2014, Sin, Annavi et al. 2015). Finally, some studies have found no evidence of MHC-dependent sexual selection and suggested it may not be a universal phenomenon (Paterson and Pemberton 1997, Westerdahl 2004, Huchard, Knapp et al. 2010, Sepil, Radersma et al. 2015, Yu, Nie et al. 2018). However, few previous studies have investigated all possible mechanisms together within the same study. A meta-analysis of MHC-dependent sexual selection in non-model species found support for female choice for MHC diversity and selection for dissimilarity when the MHC is characterized across multiple loci, but selection for particular MHC alleles was not examined (Kamiya, O’Dwyer et al. 2014). Therefore, more empirical studies are needed to draw definite conclusions about MHC-dependent sexual selection.

Future studies on MHC-dependent sexual selection would ideally include advances in a number of aspects of data quality. First, accurate estimates of relatedness should be examined at the same time as MHC-dependent sexual selection. Rather than being the actual cues for sexual selection, MHC variation could be incidentally associated with signals of genome-wide relatedness. Thus, signatures of MHC-dependent disassortative mating could be a by-product of inbreeding avoidance (Hurst, Thom et al. 2005, Sherborne, Thom et al. 2007). In the previous studies reported above (Bonneaud, Chastel et al. 2006, Setchell, Charpentier et al. 2010, Huchard, Baniel et al. 2013, Bichet, Penn et al. 2014, Winternitz, Promerova et al. 2015, Ferrandiz-Rovira, Allaine et al. 2016), relatedness was usually estimated from small panels of microsatellite markers rather than genome-wide SNPs, which may have been imprecise. Second, behavioural observations of mating are required to differentiate pre- and post-copulatory MHC-dependent sexual selection, especially since pre- and post-copulatory sexual selection on MHC genes could occur in opposite directions. For example, pre-copulatory sexual selection favouring MHC-dissimilar partners has been demonstrated in a population of salmon (*Salmo salar*) (Landry, Garant et al. 2001) while post-copulatory sexual selection favoured MHC-similar partners in another salmon study system (Yeates, Einum et al. 2009). However, few studies have examined whether MHC-dependent sexual selection occurs at the pre- and post-copulatory sexual selection stages using field observations of mating behaviour in the same population, except for a study of mouse lemur (*Microcebus murinus*)(Schwensow, Eberle et al. 2008). Third, detailed knowledge about MHC haplotype structure are needed to study MHC-dependent sexual selection. MHC loci genes are usually in strong linkage disequilibrium and inherited as haplotypes such that deviation of random mating based on MHC genes may be imprecisely measured without haplotype information. However, only a handful of studies in wild populations has managed to haplotype the MHC region (Huchard, Weill et al. 2008, Niskanen, Kennedy et al. 2014, Sin, Annavi et al. 2014, Gaigher, Burri et al. 2016) and few studies have investigated MHC-dependent sexual selection using MHC haplotypes (Sin, Annavi et al. 2015). Finally, larger sample sizes are essential in future studies to secure confident results (Kamiya, O’Dwyer et al. 2014, Hoover and Nevitt 2016, Winternitz, Abbate et al. 2017). A recent study documented the impact of sample size on error rates and effect sizes and suggested a sample size of 500 mating pairs is required for testing MHC-dependent sexual selection (Hoover and Nevitt 2016) which was not always available in previous studies.

In this study, we used a free-living population of Soay sheep (*Ovis aries*) on the island of Hirta, St Kilda to investigate MHC-dependent sexual selection. An individual-based study has been carried out on the population since 1985 (Clutton-Brock and Pemberton 2004). A large number of male-female consorts has been recorded during the rut each year and a multi-generation pedigree including most study individuals has been constructed. Previous studies have suggested intensive male-male competition and male mate choice in Soay sheep (Preston, Stevenson et al. 2003, Preston, Stevenson et al. 2005). Also, selection on load and tolerance of gastrointestinal parasites has been demonstrated (Hayward, Wilson et al. 2011, Hayward, Nussey et al. 2014). Therefore, we assume there could be MHC-dependent sexual selection to increase an offspring’s fitness to better combat parasites. However, a previous study using five MHC-linked microsatellite loci genotyped in between 887 and 1209 individuals born between 1985 and 1994 found no evidence for MHC-dependent assortative or disassortative mating in this population (Paterson and Pemberton 1997). Recently, using genotyping-by-sequencing, a total of eight MHC class II haplotypes have been identified in the study populations and a large number of individuals alive between 1989 and 2012 have been diplotyped (Dicks, Pemberton et al. 2019, Dicks, Pemberton et al. 2020). In addition, pairwise relatedness based on genome-wide SNPs is available between most individuals. This genetic and genomic data combined with a large number of consort and parentage records enabled us to test MHC-dependent sexual selection more thoroughly than before using Monte Carlo simulations. We aimed to test for specific mechanisms of MHC-dependent sexual selection by asking the following questions: 1) Are individuals carrying specific MHC haplotypes favoured during mating? 2) Are MHC-heterozygote individuals favoured during mating? 3) Is there MHC-dependent disassortative or assortative mating? 4) If there is disassortative mating, is it based on haplotype divergence? 5) Is there inbreeding preference or avoidance? 6) If any signature of non-random mating is detected, does it occur at the pre- or post-copulatory stage? 7) If there is any signature of MHC-based mating, is this signature independent of genome-wide heterozygosity or relatedness?

## Methods

### Study population and data collection

An unmanaged population of Soay sheep has resided unmanaged on the island of Hirta, St Kilda since 1932 when they were introduced there from the nearby island of Soay. From 1985, a longitudinal individual-based study has been conducted on the sheep resident in the Village Bay area of Hirta. Nearly all individual Soay sheep living in the study area have been followed from birth, through all breeding attempts, to death. Lambs born as singletons, twins or (rarely) triplets are ear tagged shortly after birth in April or May, sampled for genetic analysis and weighed. Any missed lambs or immigrant adults are captured, tagged and sampled in an August catch up or in the rut in November. The population is regularly censused throughout the year with individual locations recorded (Clutton-Brock and Pemberton 2004).

A large number of study area sheep alive since 1989 have been genotyped on the Illumina Ovine 50K SNP array. Parentage was determined for each individual using a subset of 315 unlinked SNPs derived from the SNP array using the R library Sequoia (Berenos, Ellis et al. 2014, Huisman 2017).

### Rut behavioural data

Soay sheep have a promiscuous mating system with intensive male-male competition as well as male mate choice (Preston, Stevenson et al. 2003, Preston, Stevenson et al. 2005). The onset of the rut in early November is marked by increasing male aggression as males roam and search for oestrous females across the study area. Males compete to gain access to oestrous females and the winner defends and mates with the female repeatedly over several hours in a so-called ‘consort’. However, only large and mature males with big horns can defend a female for long. Younger, smaller males and those with scurred horns constantly search for oestrous females, chase them and get quick matings if they can. Matings are mostly between males and females aged one year or older, but some male lambs aged seven months obtain matings and some female lambs give birth at the age of one (see Supplementary table 1). Throughout the rut the study area is continually monitored for consorts throughout each day, with consorts defined as being a close spatial relationship between a male and female with frequent male courtship and defence of a receptive female (Clutton-Brock and Pemberton 2004).

### Molecular data

The molecular data used in this study included MHC class II diplotypes and pairwise genomic relatedness. We used a two-step haplotyping method involving characterisation of the MHC haplotypes present and then Kompetitive Allele-specific PCR (KASP) genotyping to impute haplotypes of individuals that lived in the study area between 1989 and 2012. First, seven expressed loci (DRB1, DQA1, DQA2, DQA2-like, DQB1, DQB2 and DQB2-like) within the MHC class IIa region were characterized in 118 Soay sheep using genotyping-by-sequencing which identified eight haplotypes named A to H (Dicks, Pemberton et al. 2019). Second, a panel of 13 SNPs located in the MHC class IIa region haplotypes including 11 SNPs from the Ovine Infinium high density SNP array (Illumina) and two other SNPs located within DQA1 gene were selected to impute eight haplotypes using Kompetitive Allele-specific PCR (KASP) in 5951 Soay sheep (Dicks, Pemberton et al. 2020). After genotyping quality control and pedigree checking, 5349 individuals were successfully diplotyped. The individual inbreeding coefficients and the pairwise genomic relatedness between all individuals were calculated using GCTA (Yang, Lee et al. 2011) and DISSECT respectively (Canela-Xandri, Law et al. 2015) based on 37K polymorphic SNPs from the Ovine 50K SNP array (Illumina). The X chromosome and chromosome 20 where the MHC genes are located were excluded when calculating the pairwise genomic relatedness.

### Assortative mating analysis

We performed Monte-Carlo simulations to examine whether there were MHC-dependent or relatedness-dependent mating patterns in Soay sheep.

We first selected all females and males older than one year which were diplotyped at the MHC and genotyped on the SNP chip into a “primary mating pool”. We focused on mating between adult sheep because relatively few offspring have a juvenile parent (that is, a male lamb for a father and/or a female lamb for a mother; see Supplementary table 1) and including the large number of non-reproductive male and female lambs present each breeding season could have biased the simulations. We excluded individuals whose birth year was unknown, as they do not live mainly in our study area and we had no information about when they start to be involved in mating. For the few individuals with no recorded death year, death year was estimated from the last-seen year recorded in the census data or the last year they sired or gave birth to offspring. A total of 889 females and 821 males were included in the primary mating pool dataset and each of them is recorded once per year they were alive (Supplementary table 1).

Second, we extracted all the consorts between any male and female in the primary mating pool dataset as the “consort dataset”. Multiple observations of the same pair together on the same day were counted as one consort observation.

Third, we assembled an “observed parentage dataset” comprising all mother-father-offspring trios in which the offspring birth year was known and the parents were successfully diplotyped at the MHC and genotyped on the SNP chip. To be consistent between the primary mating pool and observed parentage dataset, we excluded trios in which either of the parents was a lamb or an adult not included in primary mating pool, such that all parents could be sampled from the primary mating pool. In total, 2068 trios were included in the observed parentage dataset (Supplementary table 2). Twins and triplets were not common in our dataset comprising less than a quarter of all individuals born. Twins are usually half-sibs with different fathers, coming from different mating events (Supplementary table 3). Thus, we treated each offspring as an independent data point. Finally, the null model of random mating (adjusted random mating model) was defined for simulation.

For each year, we first randomly assigned a male or a female living in the same year from the primary mating pool to replace each known mother and father in the observed parentage dataset and then adjusted the record of the sampled sheep based on the annual breeding success (number of offspring produced each year) of the replaced sheep to produce an “adjusted mating pool”. Then, each offspring in the observed parentage dataset was randomly assigned a father and mother in the year before its birth year without replacement from the adjusted mating pool. Annual breeding success in the simulated results was the same as the observed parentage dataset, while lifetime breeding success differed slightly from the observed parentage dataset with the mean of lifetime breeding success lower in the simulated data than that from the observed parentage dataset (Supplementary figure 1). The model was simulated for 10000 iterations in R v.3.5.2 using a custom script (R Core Team 2013).

Previous studies have suggested accounting for spatial distance when designing null models of random mating (Huchard, Baniel et al. 2013, Sepil, Radersma et al. 2015). However, during the rut, male sheep rove around the entire study area to search for oestrous females such that there is little evidence of spatially-restricted mating patterns (Clutton-Brock and Pemberton 2004). Therefore, we did not account for spatial distance in our null models.

After simulation, we summarised the results of all the iterations using various indices in response to the questions we proposed: 1) The frequency of each haplotype in simulated mothers and fathers. 2) The ratio of homozygote: heterozygote in simulated mothers and fathers. 3) The average number of shared MHC haplotypes between simulated parents and the proportion of simulated parents sharing 0, 1 and 2 haplotypes. 4) To account for MHC functional variation, the pairwise divergence of MHC haplotypes between each simulated parent pair, which has been found to be associated with fitness and parasite resistance (Wakeland, Boehme et al. 1990, Lenz, Mueller et al. 2013, Pierini and Lenz 2018), was first measured by the proportion of the amino acid sequence that differed between them (p-distance) (Henikoff 1996). We then defined two indices AAdist and distmax as the mean and maximum MHC divergence between the 4 possible haplotype combinations of each simulated parent pair respectively. Finally, the mean of AAdist and distmax were calculated. 5) The mean and median of genomic relatedness between simulated parents.

These indices were also calculated for the real data using the “consort dataset” and “observed parentage dataset”. Specifically, the first two indices were measured in mated females and males (consort dataset) or in mothers and fathers (observed parentage dataset) while the last three indices were measured between consort pairs or between genetic parents. For each index, statistical significance (*p* value) was determined by comparing the index in the real data with the 2.5% and 97.5% tails of the distribution of the index in the simulated results. To account for multiple testing, we applied a Bonferroni correction to the eight haplotype frequency tests across two sexes, and significant results were determined based on the refined critical *p*-value (*p* = 0.0015625).

Once significant results were identified, we determined whether the distortion occurred at the pre- or post-copulatory stage based on the following logic. 1) If indices in the consort dataset and observed parentage dataset both deviate from expectations of random mating, this is evidence of pre-copulatory selection. 2) If indices in the consort dataset and observed parentage dataset both deviate from expectations of random mating but in opposite directions, or only indices in the consort dataset deviated, this is evidence of both pre- and post-copulatory selection acting in different directions. 3) If only indices in the observed parentage dataset deviate from expectations of random mating, this is evidence of post-copulatory selection. 4) If indices in the consort dataset and observed parentage dataset were both in line with expectations of random mating, this suggests no evidence of sexual selection.

### Generalized linear mixed models

We used generalized linear mixed models to differentiate MHC and genomic effects on non-random mating. We drew up a matrix of all pairwise combinations of males and females in the primary mating pool in a given year. For each pair, we then recorded their consort and breeding success (0/1) based on the real data from consort dataset and observed parentage dataset respectively, with 0 meaning no successful consort or offspring observed and 1 meaning consort or offspring observed. Then, we investigated consort and breeding success as response in separate binomial regressions. In each model, year and sire ID were fitted as random effects while genome-wide heterozygosity of each mother and father measured as inbreeding coefficient, MHC heterozygosity of each mother and father, genomic relatedness and the number of shared MHC haplotypes between each pair were fitted simultaneously as fixed effects. The model was run in R v.3.5.2 using R package lme4.

## Results

### Haplotype frequency tests

The frequency of haplotype C in both consort males and fathers was significantly lower than expected under the null model, even after Bonferroni correction. In addition, the frequency of haplotype G in both consort males and fathers and the frequency of haplotype H in fathers tended to be higher than expected but these results were not significant after Bonferroni correction (Figure 1).

**Figure 1.**
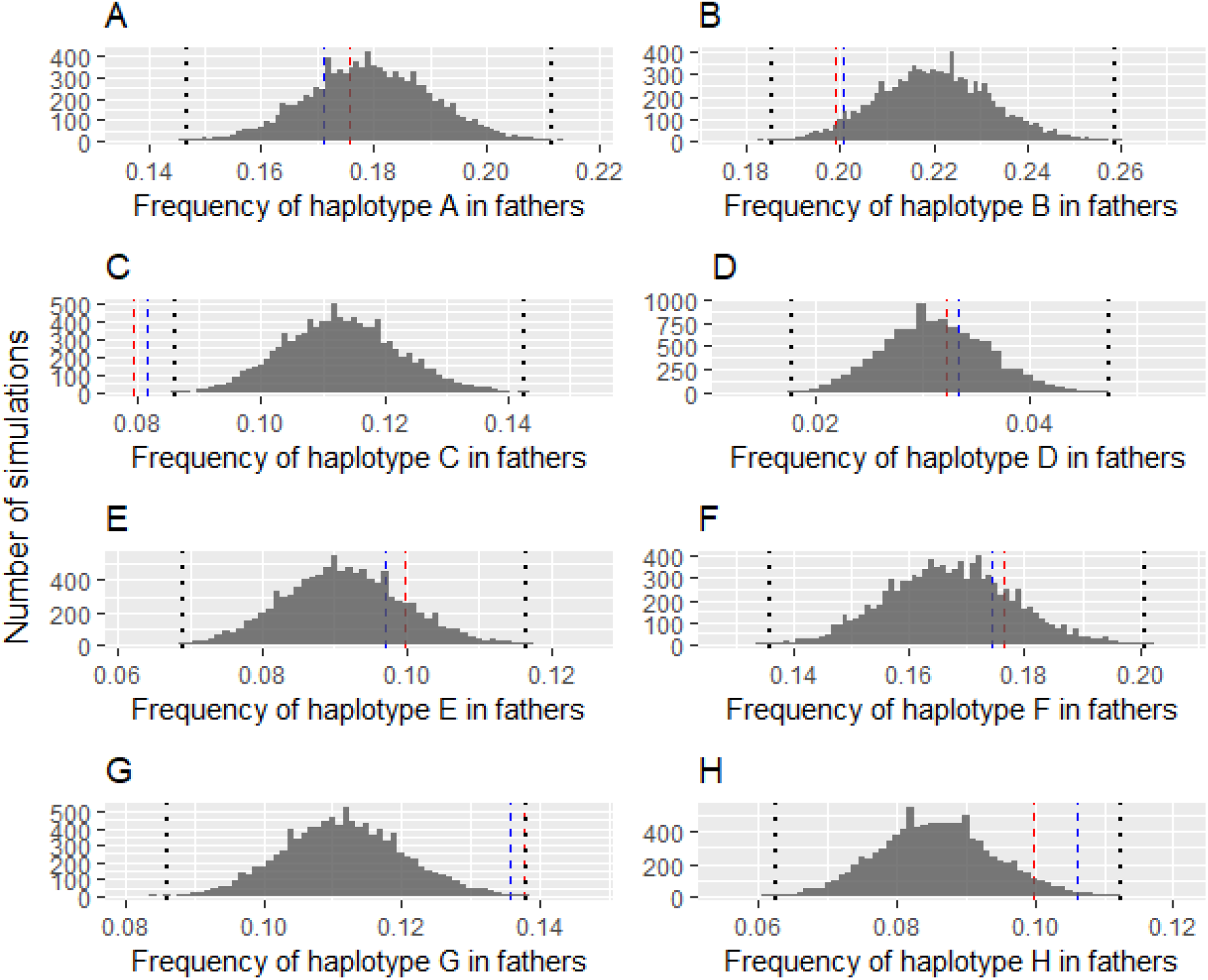
Results of MHC haplotype frequency tests in fathers following Monte Carlo simulation. Histograms represent the result of simulations with dotted black lines representing the critical p-values after Bonferroni correction. The red and blue dashed blue lines show the observed MHC haplotype frequency in the consort and observed parentage dataset respectively. Males carrying Haplotype C are rarer than expected in both the consort and parentage dataset.

We did not find any significant deviation of haplotype frequency in either consort females or mothers relative to random mating. The frequency of haplotype F and H in consort females tended to be lower and higher respectively. In addition, the frequency of haplotypes A and G in mothers tended to be lower and higher respectively. However, none of these patterns was significant after Bonferroni correction (Supplementary figure 2).

### Diplotype-based tests

Regarding individual MHC heterozygosity, we found that the ratio of homozygote to heterozygote in consort females was significantly higher than expected under the null model However, the ratio in mothers was in line with expectation under the null model (Figure 2a). We found the average number of shared haplotypes between parents was significantly lower compared with the null expectation, but this pattern was not observed in consort dataset (Figure 2c). In addition, the proportion of parents sharing 0 haplotype was significantly higher than in the simulated results while the proportion of parents sharing 1 haplotype was significantly lower than the simulated results (Figure 2d-f). However, MHC divergence measured as amino acid sequence differences between parents was in line with expectation of random mating (Figure 2g and h).

**Figure 2.**
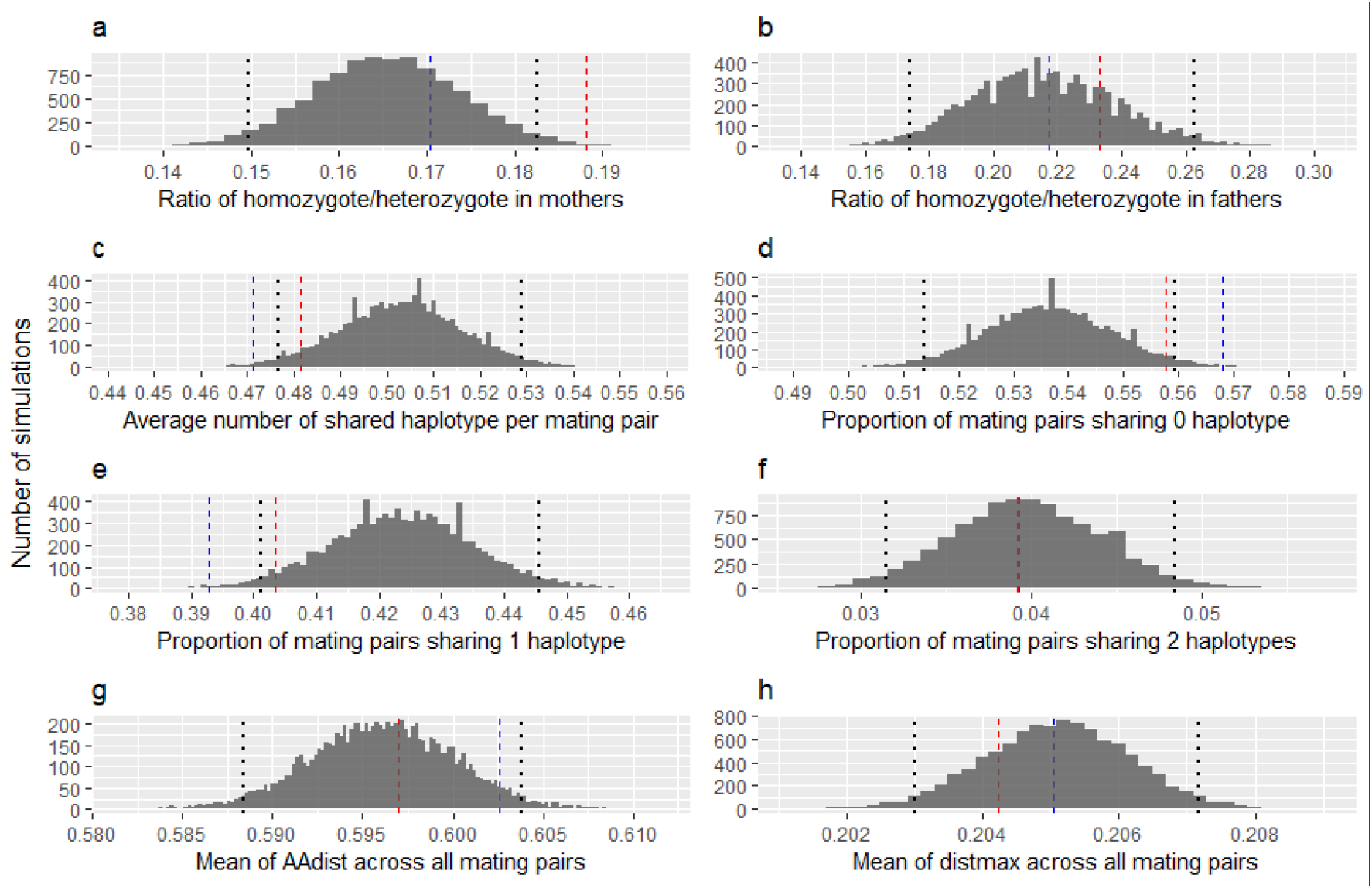
Results of MHC diplotype-based tests following Monte Carlo simulation. Histograms represent the result of simulations, with dotted lines representing the 2.5% and 97.5% tails of the distributions. The red and blue dashed blue lines show observed values for the indices in the consort and observed parentage datasets respectively. Homozygote females are commoner than expected in the consort but not the parentage dataset and parents share fewer haplotypes than expected.

### Genome-wide relatedness tests

We found mean and median genomic relatedness between consort pairs and between parents were significantly lower than expected under the null model (Figure 3).

**Figure 3.**
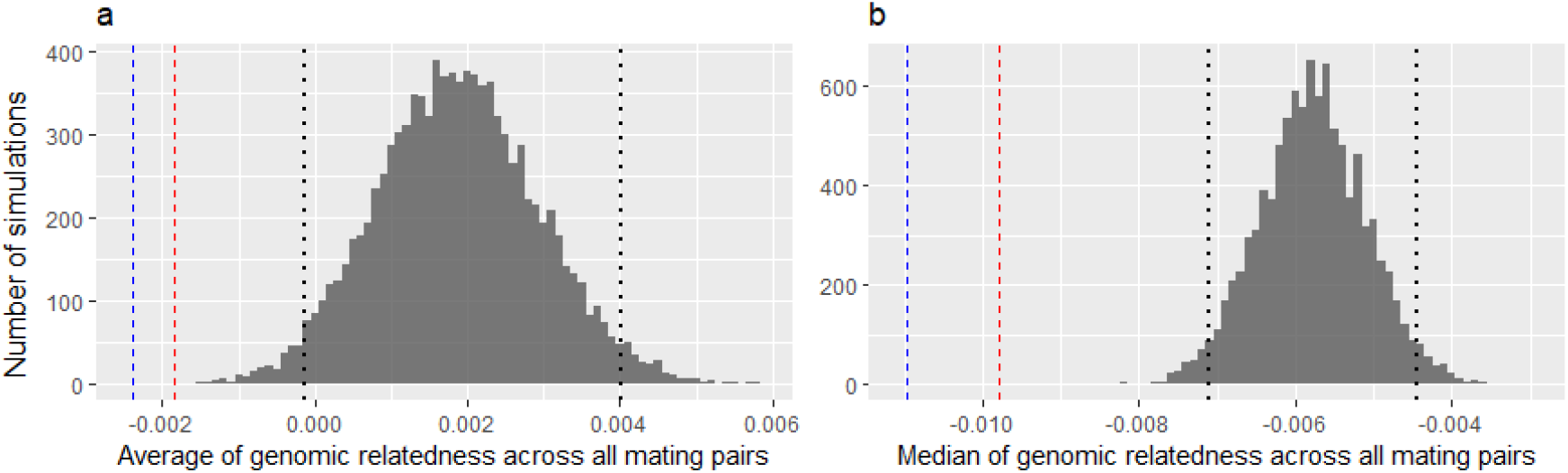
Results of tests of pairwise genomic relatedness following Monte Carlo simulation. Histograms represent the result of simulations, with dotted lines representing the 2.5% and 97.5% tails of the distributions. The red and blue dashed blue lines show observed values for genomic relatedness in the consort and observed parentage dataset respectively. Partners are less related than expected in both the consort and the parentage datasets.

### Generalized linear mixed models

We found a negative association between consort success and female MHC heterozygosity and positive associations between both consort and breeding success and male genome-wide heterozygosity measured as inbreeding coefficients. We found a negative association between both consort and breeding success and genomic relatedness, but no association between consort and breeding success and number of shared MHC haplotypes (Table 1).

**Table 1.** Results of generalized linear mixed model testing associations between consort/breeding success and MHC/genomic heterozygosity and relatedness. Significant effects and standard errors are marked with asterisk (* *p*<0.05, \*\**p*<0.01) and shown in brackets respectively.

|  | Consort Success | Breeding success |
| --- | --- | --- |
| Fixed effects |  |  |
| MHC heterozygosity of mothers | -0.154 (0.046)** | -0.032(0.063) |
| MHC heterozygosity of fathers | -0.228 (0.217) | -0.137(0.167) |
| Inbreeding coefficient of mothers | 0.335 (0.671) | -1.280(0.993) |
| Inbreeding coefficient of fathers | -12.611 (2.093)** | -10.591(3.845)** |
| Number of shared MHC haplotypes | -0.016 (0.031) | -0.064(0.041) |
| Genomic relatedness | -1.125 (0.359)** | -0.950(0.476)* |
| Random effects |  |  |
| Sire ID | 3.822(1.955) | 1.731(1.316) |
| Year | 1.248(1.117) | 0.359(0.599) |

## Discussion

In this study, we investigated MHC-dependent sexual selection in a free-living sheep population using Monte Carlo simulations. By comparing the result of simulated and real data, we examined whether there is deviation from random mating depending on MHC variation. We found haplotype C was disfavoured in comparison with random expectation in both consort males and fathers. We found no evidence that MHC heterozygotes were favoured in either sex. Instead, we found MHC homozygote females were over-represented in consort pairs but this pattern was not observed in actual mothers. We found the average number of shared MHC haplotypes between parents was lower compared with null expectation, but this pattern was not observed in the consort dataset. In addition, the proportion of parent pairs sharing no haplotype was higher and that of parent pairs sharing one haplotype was lower than expected under random mating. Finally, we found evidence of inbreeding avoidance, as the mean and median pairwise genomic relatedness in the consort and observed parentage datasets were significantly lower than expected under random mating. When fitting MHC and genomic effects in the same model of consort or parentage, we could not demonstrate an independent effect of disassortative mating based on MHC haplotype sharing, but we found the deviation towards MHC homozygote females in the consort data was independent of genome-wide heterozygosity.

In our study, sexual selection on a specific MHC haplotype (C) was probably due to differences in male competitive ability, since the frequency of C was rarer than expected, not only in male parents but also in consort males. Few previous studies have reported specific MHC variants being favoured or disfavoured during mating as the high polymorphism of MHC genes requires a large sample size to detect sexual selection on specific MHC variants (Eizaguirre, Yeates et al. 2009). In this study, our finding for haplotype C in male parents was consistent with a negative association between MHC haplotype C and male breeding success found in a recent study on MHC-fitness associations (Huang, Dicks et al. 2020). The fact that haplotype C males are also less often observed in consort than expected indicates that the effect of haplotype C is expressed at the pre-copulatory rather than post-copulatory stage.

Our finding that females observed in consorts are more homozygous than expected, an effect which is opposite to expectation, is not found in the observed parentage data and is independent of genome-wide heterozygosity, is puzzling and requires explanation. One hypothesis is that MHC-homozygous females are less likely to conceive in a given oestrus cycle and therefore return to oestrus 14 days later. This in turn would enrich our consort data set for such females. Alternatively, if homozygous females are less attractive in some way, perhaps they are less likely to be in long, stable consorts and instead experience multiple short consorts, which would again enrich the consort data set for homozygous females. These possibilities require further investigation within our dataset but are beyond the scope of this paper.

At first sight, our results also suggest that there is sexual selection based on MHC compatibility, but our tests suggest this effect is not independent of an inbreeding avoidance effect. In a population with limited dispersal and severe inbreeding depression, inbreeding avoidance through kin recognition could arise to reduce the cost of inbreeding (Pusey and Wolf 1996, Szulkin, Stopher et al. 2013, Duthie and Reid 2016). Previous studies have proposed MHC variation could be used as a cue for inbreeding avoidance and MHC-associated odour variation has been reported in a wide range of taxa including fish (Olsen, Grahn et al. 1998, Milinski, Griffiths et al. 2005), reptiles (Olsson, Madsen et al. 2003), birds (Leclaire, Strandh et al. 2017) and mammals (Wedekind, Seebeck et al. 1995, Wedekind and Furi 1997, Yamazaki, Beauchamp et al. 2000, Roberts, Gosling et al. 2008). However, if MHC haplotype sharing is associated with relatedness, as we have shown in Soay sheep, MHC-disassortative mating could be a by-product of inbreeding avoidance. Few studies have been able to test this hypothesis, however a study of grey mouse lemur revealed both inbreeding avoidance and MHC-dependent disassortative mating, and suggested that observed deviations from random mating at the MHC are driven by the functionally important MHC gene DRB rather than resulting passively from inbreeding avoidance. In that study, MHC-dependent disassortative mating was detected at the DRB locus only for amino acid sequence and functional similarity rather than number of shared MHC alleles (Huchard, Baniel et al. 2013). In contrast, when studied in terms of MHC divergence measured as amino acid sequence differences between parents in Soay sheep, our results were in line with the expectation from random mating (Figure 2g and h). In Soay sheep, inbreeding depression has been documented repeatedly using different approaches (Coltman, Pilkington et al. 1999, Overall, Byrne et al. 2005, Berenos, Ellis et al. 2016, Stoffel, Johnston et al. 2020), so it is possible the sheep have evolved inbreeding avoidance. Using genomic relatedness calculated from a large number of SNPs, we demonstrated genomic inbreeding avoidance in Soays at the pre-copulatory stage for the first time. When tested in the same model, MHC haplotype sharing was not significant. We therefore cannot claim that the apparent disassortative mating based on haplotype sharing is anything but a correlated effect of inbreeding avoidance.

Our results differ from a previous study in Soay sheep (Paterson and Pemberton 1997) which found no evidence for MHC-dependent assortative or disassortative mating. In the current study, we found evidence of MHC-dependent disassortative mating, at the post-copulatory stage which carried through to the parentage stage, based on the number of shared MHC haplotypes. Reasons for this difference include the fact our data consisted of MHC class II haplotypes rather than MHC-linked microsatellite markers, and these two approaches do not have a perfect read through. Also, our sample sizes are very much larger and our methodology (Monte Carlo simulation) is different from the previous study which used a likelihood-based approach.

By using a large number of consort observations, we were able to differentiate MHC-dependent sexual selection at the pre- and post-copulatory stages. In this area, field observations were first used in a study of mouse lemur which demonstrated post-copulatary MHC-dependent disassortative mating (Schwensow, Eberle et al. 2008). Here, we used both the consort and the observed parentage datasets to examine MHC-dependent sexual selection. We found sexual selection against a specific MHC haplotype at pre-copulatory stage and sexual selection favouring MHC compatibility at the post-copulatory stage. Interestingly, we found that the ratio of homozygote:heterozygote was significantly higher in consort females than the simulated results but this pattern were not observed in mothers. These results indicate the value of using field observations to differentiate pre- and post-copulatory sexual selection.

In this study, we examined whether there was MHC-dependent sexual selection in a population of free-living Soay sheep. Benefiting from intermediate MHC polymorphism, high quality genetic and genomic information, intensive field observations and large sample size, we have demonstrated sexual selection based on a specific MHC haplotype at the pre-copulatory stage and MHC compatibility at the post-copulatory stage occurs simultaneously. We have also demonstrated sexual selection against female MHC heterozygosity in dependent of genome-wide heterozygosity during the rut. Finally, we report inbreeding avoidance in this population for the first time and find that we cannot show an independent effect of disassortative mating based on the MHC. Our results suggest that multiple mechanisms of MHC-dependent sexual selection could act simultaneously in Soay sheep and that it is necessary to have an exhaustive examination of all possible mechanisms when investigating MHC-dependent sexual selection.

## Acknowledgements

We thank the National Trust for Scotland for permission to work on St. Kilda and QinetiQ, Eurest and Kilda Cruises for logistics and support. We thank I. Stevenson and many volunteers who have collected field data and samples, especially in the rut, and all those who have contributed to keeping the project going. The MHC diplotyping method and the diplotype data set were developed and generated by Kara Dicks and Susan Johnston. SNP genotyping was conducted at the Wellcome Trust Clinical Research Facility Genetics Core. Field data collection has been supported by NERC (NE/M002896/1) over many years, the diplotyping was supported by the BBSRC and Royal Society and most of the SNP genotyping was supported by the European Research Council (AdG 250098). Wei Huang is supported by Edinburgh Global research scholarship.

## Data Accessibility and Benefit-Sharing Statement

All the data and R script pf this manuscript are available through the following link: https://figshare.com/articles/dataset/MHC-sexual_selection-St_Kilda_Soay_sheep/13277081

## Author contribution

W.H and J.M.P designed the study. J.G.P conducted the field observations. W.H analysed the data and wrote the manuscript. All the authors contributed to the final version of the manuscript.

## Notes

### Competing Interest Statement

The authors have declared no competing interest.

## References

Berenos, C., P. A. Ellis, J. G. Pilkington and J. M. Pemberton (2014). “Estimating quantitative genetic parameters in wild populations: a comparison of pedigree and genomic approaches.” Molecular Ecology 23(14): 3434–3451.

Berenos, C., P. A. Ellis, J. G. Pilkington and J. M. Pemberton (2016). “Genomic analysis reveals depression due to both individual and maternal inbreeding in a free-living mammal population.” Molecular Ecology 25(13): 3152–3168.

Bichet, C., D. J. Penn, Y. Moodley, L. Dunoyer, E. Cellier-Holzem, M. Belvalette, A. Gregoire, S. Garnier and G. Sorci (2014). “Females tend to prefer genetically similar mates in an island population of house sparrows.” Bmc Evolutionary Biology 14.

Bonneaud, C., O. Chastel, P. Federici, H. Westerdahl and G. Sorci (2006). “Complex Mhc-based mate choice in a wild passerine.” Proceedings of the Royal Society B-Biological Sciences 273(1590): 1111–1116.

Canela-Xandri, O., A. Law, A. Gray, J. A. Woolliams and A. Tenesa (2015). “A new tool called DISSECT for analysing large genomic data sets using a Big Data approach.” Nature Communications 6.

Clutton-Brock, T. H. and J. M. Pemberton (2004). Soay sheep: dynamics and selection in an island population, Cambridge University Press.

Coltman, D. W., J. G. Pilkington, J. A. Smith and J. M. Pemberton (1999). “Parasite-mediated selection against inbred Soay sheep in a free-living, island population.” Evolution 53(4): 1259–1267.

Cutrera, A. P., M. S. Fanjul and R. R. Zenuto (2012). “Females prefer good genes: MHC-associated mate choice in wild and captive tuco-tucos.” Animal Behaviour 83(3): 847–856.

Dicks, K. L., J. M. Pemberton and K. T. Ballingall (2019). “Characterisation of major histocompatibility complex class IIa haplotypes in an island sheep population.” Immunogenetics 71(5-6): 383–393.

Dicks, K. L., J. M. Pemberton, K. T. Ballingall and S. E. Johnston (2020). “Haplotyping MHC class Ila by high throughput screening in an isolated sheep population.” bioRxiv: 2020.2007.2020.212225.

Duthie, A. B. and J. M. Reid (2016). “Evolution of Inbreeding Avoidance and Inbreeding Preference through Mate Choice among Interacting Relatives.” American Naturalist 188(6): 651–667.

Eizaguirre, C., S. E. Yeates, T. L. Lenz, M. Kalbe and M. Milinski (2009). “MHC-based mate choice combines good genes and maintenance of MHC polymorphism.” Molecular Ecology 18(15): 3316–3329.

Ejsmond, M. J., J. Radwan and A. B. Wilson (2014). “Sexual selection and the evolutionary dynamics of the major histocompatibility complex.” Proceedings of the Royal Society B-Biological Sciences 281(1796).

Ekblom, R., S. A. Saether, M. Grahn, P. Fiske, J. A. Kalas and J. Hoglund (2004). “Major histocompatibility complex variation and mate choice in a lekking bird, the great snipe (Gallinago media).” Molecular Ecology 13(12): 3821–3828.

Ferrandiz-Rovira, M., D. Allaine, M. P. Callait-Cardinal and A. Cohas (2016). “Mate choice for neutral and MHC genetic characteristics in Alpine marmots: different targets in different contexts?” Ecology and Evolution 6(13): 4243–4257.

Gaigher, A., R. Burri, W. H. Gharib, P. Taberlet, A. Roulin and L. Fumagalli (2016). “Family-assisted inference of the genetic architecture of major histocompatibility complex variation.” Mol Ecol Resour 16(6): 1353–1364.

Griggio, M., C. Biard, D. J. Penn and H. Hoi (2011). “Female house sparrows “count on” male genes: experimental evidence for MHC-dependent mate preference in birds.” Bmc Evolutionary Biology 11.

Grob, B., L. A. Knapp, R. D. Martin and G. Anzenberger (1998). “The major histocompatibility complex and mate choice: Inbreeding avoidance and selection of good genes.” Experimental and Clinical Immunogenetics 15(3): 119–129.

Han, Q. H., R. N. Sun, H. Q. Yang, Z. W. Wang, Q. H. Wan and S. G. Fang (2019). “MHC class I diversity predicts non-random mating in Chinese alligators (Alligator sinensis).” Heredity 122(6): 809–818.

Hayward, A. D., D. H. Nussey, A. J. Wilson, C. Berenos, J. G. Pilkington, K. A. Watt, J. M. Pemberton and A. L. Graham (2014). “Natural Selection on Individual Variation in Tolerance of Gastrointestinal Nematode Infection.” Plos Biology 12(7).

Hayward, A. D., A. J. Wilson, J. G. Pilkington, T. H. Clutton-Brock, J. M. Pemberton and L. E. B. Kruuk (2011). “Natural selection on a measure of parasite resistance varies across ages and environmental conditions in a wild mammal.” Journal of Evolutionary Biology 24(8): 1664–1676.

Henikoff, S. (1996). “Scores for sequence searches and alignments.” Current Opinion in Structural Biology 6(3): 353–360.

Hoover, B., M. Alcaide, S. Jennings, S. Y. W. Sin, S. V. Edwards and G. A. Nevitt (2018). “Ecology can inform genetics: Disassortative mating contributes to MHC polymorphism in Leach’s storm-petrels (Oceanodroma leucorhoa).” Mol Ecol.

Hoover, B. and G. Nevitt (2016). “Modeling the Importance of Sample Size in Relation to Error in MHC-Based Mate-Choice Studies on Natural Populations.” Integrative and Comparative Biology 56(5): 925–933.

Huang, W., K. L. Dicks, J. D. Hadfield, S. E. Johnston, K. T. Ballingall and J. M. Pemberton (2020). “A rare MHC haplotype confers selective advantage in a free-living ruminant.” bioRxiv: 2020.2003.2025.008565.

Huchard, E., A. Baniel, S. Schliehe-Diecks and P. M. Kappeler (2013). “MHC-disassortative mate choice and inbreeding avoidance in a solitary primate.” Molecular Ecology 22(15): 4071–4086.

Huchard, E., L. A. Knapp, J. Wang, M. Raymond and G. Cowlishaw (2010). “MHC, mate choice and heterozygote advantage in a wild social primate.” Mol Ecol 19(12): 2545–2561.

Huchard, E., M. Weill, G. Cowlishaw, M. Raymond and L. A. Knapp (2008). “Polymorphism, haplotype composition, and selection in the Mhc-DRB of wild baboons.” Immunogenetics 60(10): 585–598.

Huisman, J. (2017). “Pedigree reconstruction from SNP data: parentage assignment, sibship clustering and beyond.” Molecular Ecology Resources 17(5): 1009–1024.

Hurst, J. L., M. D. Thom, C. M. Nevison, R. E. Humphries and R. J. Beynon (2005). “MHC odours are not required or sufficient for recognition of individual scent owners.” Proceedings of the Royal Society B-Biological Sciences 272(1564): 715–724.

Jordan, W. C. and M. W. Bruford (1998). “New perspectives on mate choice and the MHC.” Heredity 81: 239–245.

Kamiya, T., K. O’Dwyer, H. Westerdahl, A. Senior and S. Nakagawa (2014). “A quantitative review of MHC-based mating preference: the role of diversity and dissimilarity.” Molecular Ecology 23(21): 5151–5163.

Landry, C., D. Garant, P. Duchesne and L. Bernatchez (2001). “‘Good genes as heterozygosity’: the major histocompatibility complex and mate choice in Atlantic salmon (Salmo salar).” Proceedings of the Royal Society B-Biological Sciences 268(1473): 1279–1285.

Leclaire, S., M. Strandh, J. Mardon, H. Westerdahl and F. Bonadonna (2017). “Odour-based discrimination of similarity at the major histocompatibility complex in birds.” Proceedings of the Royal Society B-Biological Sciences 284(1846).

Lenz, T. L., B. Mueller, F. Trillmich and J. B. W. Wolf (2013). “Divergent allele advantage at MHC-DRB through direct and maternal genotypic effects and its consequences for allele pool composition and mating.” Proceedings of the Royal Society B-Biological Sciences 280(1762).

Migalska, M., A. Sebastian and J. Radwan (2019). “Major histocompatibility complex class I diversity limits the repertoire of T cell receptors.” Proc Natl Acad Sci U S A 116(11): 5021–5026.

Milinski, M. (2006). “The major histocompatibility complex, sexual selection, and mate choice.” Annual Review of Ecology Evolution and Systematics 37: 159–186.

Milinski, M., S. Griffiths, K. M. Wegner, T. B. H. Reusch, A. Haas-Assenbaum and T. Boehm (2005). “Mate choice decisions of stickleback females predictably modified by MHC peptide ligands.” Proceedings of the National Academy of Sciences of the United States of America 102(12): 4414–4418.

Niskanen, A. K., L. J. Kennedy, M. Ruokonen, I. Kojola, H. Lohi, M. Isomursu, E. Jansson, T. Pyhajarvi and J. Aspi (2014). “Balancing selection and heterozygote advantage in major histocompatibility complex loci of the bottlenecked Finnish wolf population.” Mol Ecol 23(4): 875–889.

Olsen, K. H., M. Grahn, J. Lohm and A. Langefors (1998). “MHC and kin discrimination in juvenile Arctic charr, Salvelinus alpinus (L.).” Animal Behaviour 56: 319–327.

Olsson, M., T. Madsen, J. Nordby, E. Wapstra, B. Ujvari and H. Wittsell (2003). “Major histocompatibility complex and mate choice in sand lizards.” Proceedings of the Royal Society B-Biological Sciences 270: S254–S256.

Overall, A. D. J., K. A. Byrne, J. G. Pilkington and J. M. Pemberton (2005). “Heterozygosity, inbreeding and neonatal traits in Soay sheep on St Kilda.” Molecular Ecology 14(11): 3383–3393.

Paterson, S. and J. M. Pemberton (1997). “No evidence for major histocompatibility complex-dependent mating patterns in a free-living ruminant population.” Proceedings of the Royal Society B-Biological Sciences 264(1389): 1813–1819.

Penn, D. J. (2002). “The scent of genetic compatibility: Sexual selection and the major histocompatibility complex.” Ethology 108(1): 1–21.

Pierini, F. and T. L. Lenz (2018). “Divergent Allele Advantage at Human MHC Genes: Signatures of Past and Ongoing Selection.” Mol Biol Evol 35(9): 2145–2158.

Preston, B. T., I. R. Stevenson, J. M. Pemberton, D. W. Coltman and K. Wilson (2003). “Overt and covert competition in a promiscuous mammal: the importance of weaponry and testes size to male reproductive success.” Proceedings of the Royal Society B-Biological Sciences 270(1515): 633–640.

Preston, B. T., I. R. Stevenson, J. M. Pemberton, D. W. Coltman and K. Wilson (2005). “Male mate choice influences female promiscuity in Soay sheep.” Proceedings of the Royal Society B-Biological Sciences 272(1561): 365–373.

Pusey, A. and M. Wolf (1996). “Inbreeding avoidance in animals.” Trends in Ecology & Evolution 11(5): 201–206.

R Core Team (2013). “R: A language and environment for statistical computing..”

Radwan, J., W. Babik, J. Kaufman, T. L. Lenz and J. Winternitz (2020). “Advances in the Evolutionary Understanding of MHC Polymorphism.” Trends in Genetics 36(4): 298–311.

Rekdal, S. L., J. A. Anmarkrud, J. T. Lifjeld and A. Johnsen (2019). “Extra-pair mating in a passerine bird with highly duplicated major histocompatibility complex class II: Preference for the golden mean.” Molecular Ecology 28(23): 5133–5144.

Reusch, T. B. H., M. A. Haberli, P. B. Aeschlimann and M. Milinski (2001). “Female sticklebacks count alleles in a strategy of sexual selection explaining MHC polymorphism.” Nature 414(6861): 300–302.

Richardson, D. S., J. Komdeur, T. Burke and T. von Schantz (2005). “MHC-based patterns of social and extra-pair mate choice in the Seychelles warbler.” Proceedings of the Royal Society B-Biological Sciences 272(1564): 759–767.

Roberts, S. C., L. M. Gosling, V. Carter and M. Petrie (2008). “MHC-correlated odour preferences in humans and the use of oral contraceptives.” Proceedings of the Royal Society B-Biological Sciences 275(1652): 2715–2722.

Schwensow, N., M. Eberle and S. Sommer (2008). “Compatibility counts: MHC-associated mate choice in a wild promiscuous primate.” Proceedings of the Royal Society B-Biological Sciences 275(1634): 555–564.

Sepil, I., R. Radersma, A. W. Santure, I. De Cauwer, J. Slate and B. C. Sheldon (2015). “No evidence for MHC class l-based disassortative mating in a wild population of great tits.” Journal of Evolutionary Biology 28(3): 642–654.

Setchell, J. M., M. J. E. Charpentier, K. M. Abbott, E. J. Wickings and L. A. Knapp (2010). “Opposites attract: MHC-associated mate choice in a polygynous primate.” Journal of Evolutionary Biology 23(1): 136–148.

Sherborne, A. L., M. D. Thom, S. Paterson, F. Jury, W. E. R. Ollier, P. Stockley, R. J. Beynon and J. L. Hurst (2007). “The genetic basis of inbreeding avoidance in house mice.” Current Biology 17(23): 2061–2066.

Sin, Y. W., G. Annavi, H. L. Dugdale, C. Newman, T. Burke and D. W. MacDonald (2014). “Pathogen burden, co-infection and major histocompatibility complex variability in the European badger (Meles meles).” Mol Ecol 23(20): 5072–5088.

Sin, Y. W., G. Annavi, C. Newman, C. Buesching, T. Burke, D. W. Macdonald and H. L. Dugdale (2015). “MHC class II-assortative mate choice in European badgers (Meles meles).” Molecular Ecology 24(12): 3138–3150.

Sin, Y. W., G. Annavi, C. Newman, C. Buesching, T. Burke, D. W. Macdonald and H. L. Dugdale (2015). “MHC class II-assortative mate choice in European badgers (Meles meles).” Mol Ecol 24(12): 3138–3150.

Stoffel, M. A., S. E. Johnston, J. G. Pilkington and J. M. Pemberton (2020). “Genetic architecture and lifetime dynamics of inbreeding depression in a wild mammal.” bioRxiv.

Szulkin, M., K. V. Stopher, J. M. Pemberton and J. M. Reid (2013). “Inbreeding avoidance, tolerance, or preference in animals?” Trends in Ecology & Evolution 28(4): 205–211.

Wakeland, E. K., S. Boehme, J. X. She, C. C. Lu, R. A. McIndoe, I. Cheng, Y. Ye and W. K. Potts (1990). “Ancestral polymorphisms of MHC class II genes: divergent allele advantage.” Immunol Res 9(2): 115–122.

Wedekind, C. and S. Furi (1997). “Body odour preferences in men and women: do they aim for specific MHC combinations or simply heterozygosity?” Proceedings of the Royal Society B-Biological Sciences 264(1387): 1471–1479.

Wedekind, C., T. Seebeck, F. Bettens and A. J. Paepke (1995). “MHC-dependent mate preferences in humans.” Proc Biol Sci 260(1359): 245–249.

Westerdahl, H. (2004). “No evidence of an MHC-based female mating preference in great reed warblers.” Molecular Ecology 13(8): 2465–2470.

Winternitz, J., J. L. Abbate, E. Huchard, J. Havlicek and L. Z. Garamszegi (2017). “Patterns of MHC-dependent mate selection in humans and nonhuman primates: a meta-analysis.” Mol Ecol 26(2): 668–688.

Winternitz, J. C., S. G. Minchey, L. Z. Garamszegi, S. Huang, P. R. Stephens and S. Altizer (2013). “Sexual selection explains more functional variation in the mammalian major histocompatibility complex than parasitism.” Proceedings of the Royal Society B-Biological Sciences 280(1769).

Winternitz, J. C., M. Promerova, R. Polakova, M. Vinkler, J. Schnitzer, P. Munclinger, W. Babik, J. Radwan, J. Bryja and T. Albrecht (2015). “Effects of heterozygosity and MHC diversity on patterns of extra-pair paternity in the socially monogamous scarlet rosefinch.” Behavioral Ecology and Sociobiology 69(3): 459–469.

Yamazaki, K., G. K. Beauchamp, M. Curran, J. Bard and E. A. Boyse (2000). “Parent-progeny recognition as a function of MHC odortype identity.” Proceedings of the National Academy of Sciences of the United States of America 97(19): 10500–10502.

Yang, J., S. H. Lee, M. E. Goddard and P. M. Visscher (2011). “GCTA: a tool for genome-wide complex trait analysis.” Am J Hum Genet 88(1): 76–82.

Yeates, S. E., S. Einum, I. A. Fleming, H. J. Megens, R. J. M. Stet, K. Hindar, W. V. Holt, K. J. W. Van Look and M. J. G. Gage (2009). “Atlantic salmon eggs favour sperm in competition that have similar major histocompatibility alleles.” Proceedings of the Royal Society B-Biological Sciences 276(1656): 559–566.

Yu, L. J., Y. G. Nie, L. Yan, Y. B. Hu and F. W. Wei (2018). “No evidence for MHC-based mate choice in wild giant pandas.” Ecology and Evolution 8(17): 8642–8651.

