## Supplementary Material for "Patterns of MHC-dependent sexual selection in a free-living population of sheep"

**Table of Contents:**

| **Offspring with juvenile parents in the current study** | Page 2 |
| --- | --- |
| **Sample size of the current study** | Page 3 |
| **Twins and triplets in the current study** | Page 4 |
| **Annual and lifetime breeding success of mothers**  **and fathers in real data and the simulated results** | Page 5-6 |
| **Results of adjusted random mating model** | Page 7 |
| **Results of primary and restricted random mating model** | Page 8-12 |

**Offspring with juvenile parents in the current study**

Supplementary table 1. Sample sizes of offspring with only juvenile father or mother, both juvenile and adult parents in the observed parentage data.

| Birthyear | Juvenile father only | Juvenile mother only | Both parents juveniles | Both parents adults |
| --- | --- | --- | --- | --- |
| 1989 | 0 | 0 | 0 | 22 |
| 1990 | 5 | 8 | 4 | 36 |
| 1991 | 8 | 15 | 2 | 53 |
| 1992 | 5 | 14 | 1 | 70 |
| 1993 | 7 | 20 | 2 | 91 |
| 1994 | 4 | 10 | 0 | 73 |
| 1995 | 2 | 1 | 0 | 87 |
| 1996 | 7 | 13 | 6 | 59 |
| 1997 | 4 | 4 | 0 | 89 |
| 1998 | 1 | 4 | 0 | 120 |
| 1999 | 1 | 2 | 0 | 61 |
| 2000 | 6 | 11 | 4 | 57 |
| 2001 | 6 | 6 | 1 | 51 |
| 2002 | 2 | 0 | 0 | 66 |
| 2003 | 12 | 16 | 4 | 102 |
| 2004 | 9 | 41 | 0 | 138 |
| 2005 | 3 | 22 | 0 | 121 |
| 2006 | 4 | 13 | 0 | 129 |
| 2007 | 9 | 21 | 0 | 109 |
| 2008 | 7 | 21 | 2 | 124 |
| 2009 | 2 | 23 | 1 | 149 |
| 2010 | 0 | 6 | 1 | 7 |
| 2011 | 0 | 0 | 0 | 160 |
| 2012 | 0 | 8 | 0 | 93 |

**Sample size of the current study**

Supplementary table 2. Sample sizes of the observed parentage dataset (= the number of offspring produced) and the primary mating pool in each year of the study with number of individuals recorded at least once in the consort dataset (shown in brackets).

| Rut Year, t | Number of offspring produced in year t+1 | Number of females in the primary mating pool in year t | Number of males in the primary mating pool in year t |
| --- | --- | --- | --- |
| 1988 | 22 | 121 (63) | 56 (30) |
| 1989 | 36 | 90 (37) | 13 (11) |
| 1990 | 53 | 115 (NA) | 42 (NA) |
| 1991 | 70 | 154 (59) | 85 (31) |
| 1992 | 91 | 143 (80) | 66 (35) |
| 1993 | 73 | 172 (92) | 95 (54) |
| 1994 | 87 | 197 (111) | 101 (57) |
| 1995 | 59 | 182 (69) | 65 (33) |
| 1996 | 89 | 228 (77) | 123 (44) |
| 1997 | 120 | 238 (80) | 122 (62) |
| 1998 | 61 | 241 (101) | 120 (56) |
| 1999 | 57 | 158 (86) | 47 (24) |
| 2000 | 51 | 193 (71) | 97 (30) |
| 2001 | 66 | 247 (127) | 153 (60) |
| 2002 | 102 | 152 (74) | 63 (31) |
| 2003 | 138 | 186 (105) | 123 (40) |
| 2004 | 121 | 249 (123) | 205 (72) |
| 2005 | 129 | 174 (78) | 96 (49) |
| 2006 | 109 | 199 (88) | 120 (52) |
| 2007 | 124 | 184 (88) | 97 (47) |
| 2008 | 149 | 213 (111) | 137 (60) |
| 2009 | 7 | 234 (153) | 153 (77) |
| 2010 | 160 | 256 (99) | 149 (59) |
| 2011 | 93 | 231 (118) | 138 (58) |

**Twins and triplets in the current study**

Supplementary table 3. Sample sizes of twin and triplet offspring in the observed parentage dataset.

| Birth year | Number of genotyped individuals born as singletons or where only one of a pair of twins or set of triplets genotyped | Number of individuals born as twins with both twins genotyped | Number of individuals born as triplets and with two or more members genotyped | Number of individuals born as full-sib twins or triplets where two or more members genotyped |
| --- | --- | --- | --- | --- |
| 1988 | 15 | 7 | 0 | 4 |
| 1989 | 23 | 13 | 0 | 8 |
| 1990 | 34 | 19 | 0 | 4 |
| 1991 | 51 | 19 | 0 | 6 |
| 1992 | 47 | 44 | 0 | 26 |
| 1993 | 55 | 17 | 1 | 8 |
| 1994 | 69 | 18 | 0 | 8 |
| 1995 | 36 | 23 | 0 | 4 |
| 1996 | 71 | 18 | 0 | 4 |
| 1997 | 92 | 28 | 0 | 8 |
| 1998 | 56 | 5 | 0 | 0 |
| 1999 | 44 | 13 | 0 | 2 |
| 2000 | 38 | 13 | 0 | 2 |
| 2001 | 62 | 4 | 0 | 2 |
| 2002 | 67 | 35 | 0 | 16 |
| 2003 | 96 | 42 | 0 | 18 |
| 2004 | 104 | 17 | 2 | 12 |
| 2005 | 98 | 31 | 0 | 6 |
| 2006 | 75 | 34 | 0 | 8 |
| 2007 | 90 | 34 | 0 | 16 |
| 2008 | 123 | 26 | 0 | 12 |
| 2009 | 6 | 1 | 0 | 0 |
| 2010 | 140 | 20 | 0 | 6 |
| 2011 | 84 | 9 | 0 | 2 |

**Annual and lifetime breeding success of mothers and fathers in real data and the simulated results**

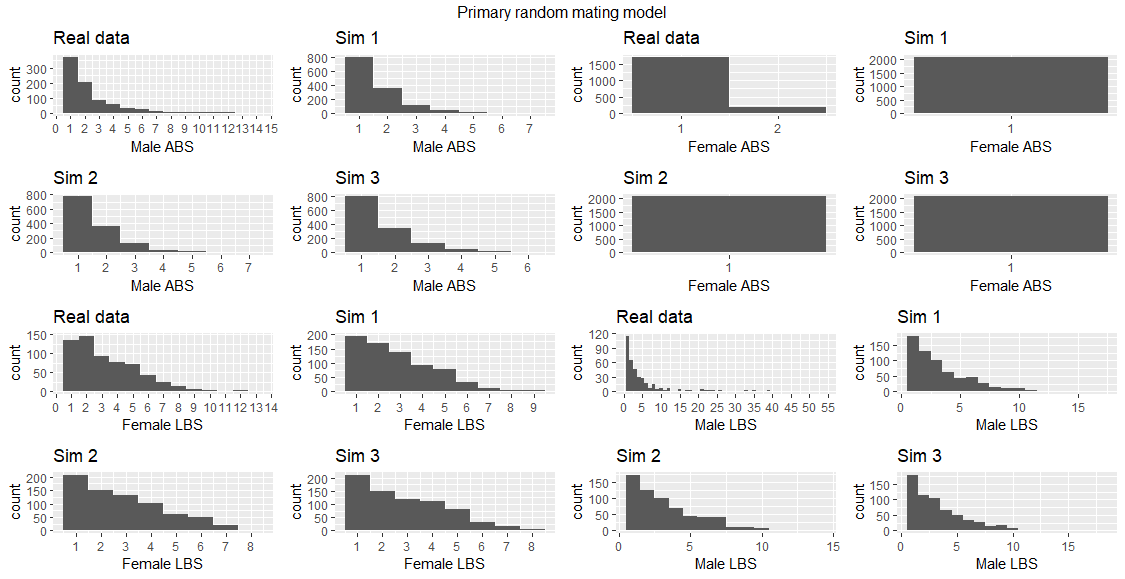

Supplementary figure 1. Annual and lifetime breeding success of mothers and fathers in the observed parentage dataset and three iterations of the simulated results under the primary random mating model.

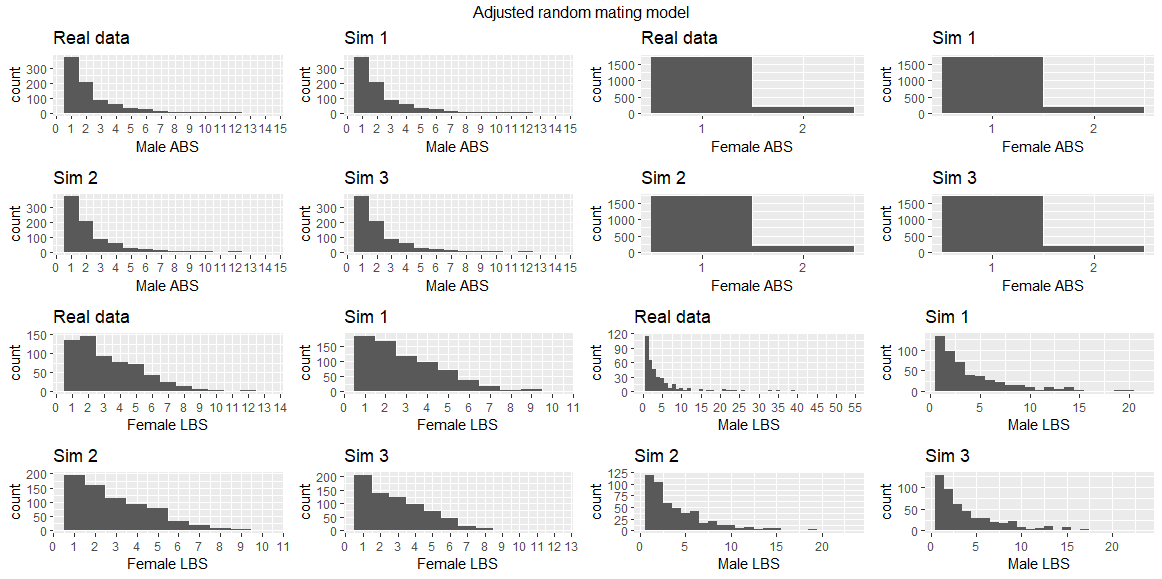

Supplementary figure 2. Annual and lifetime breeding success of mothers and fathers in the observed parentage dataset and three iterations of the simulated results under the adjusted random mating model.

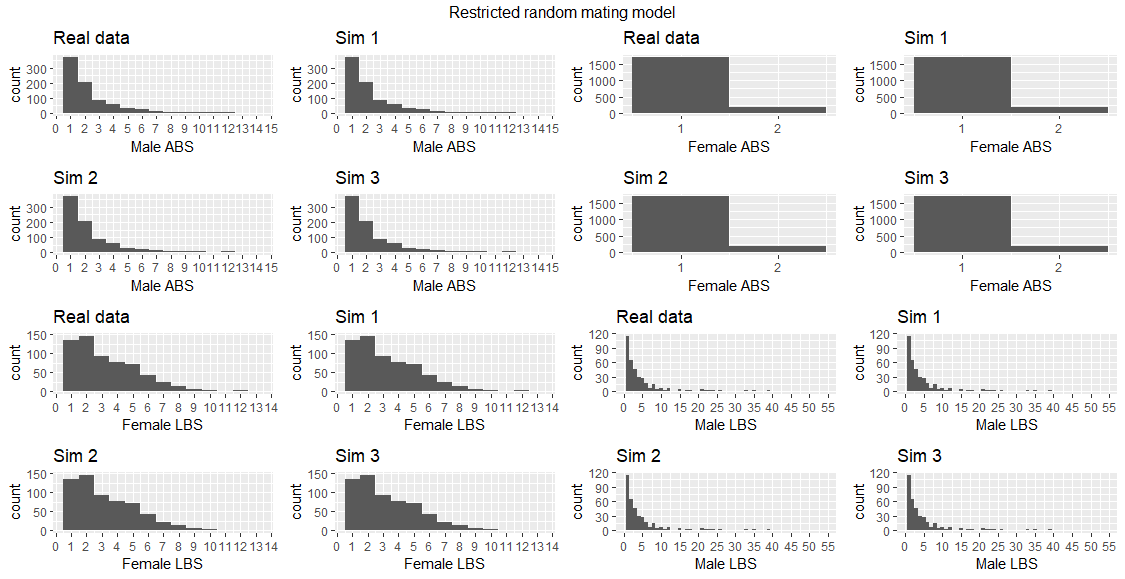

Supplementary figure 3. Annual and lifetime breeding success of mothers and fathers in the observed parentage dataset and three iterations of the simulated results under the restricted random mating model.

**Results of adjusted random mating model**

Here we show the figures for results not shown in the main paper.

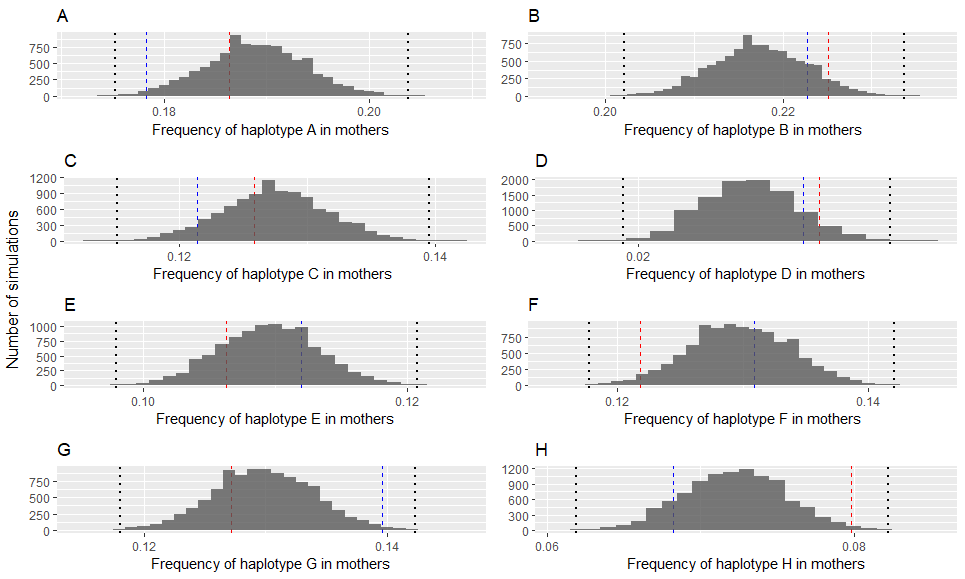

Supplementary figure 4. Results of MHC haplotype frequency in mothers under adjusted random mating model. Histograms represent the result of simulations with dotted lines represent critical p-value of the distributions after Bonferroni correction. The red and blue dashed blue lines show MHC haplotype frequency in the consort and observed parentage dataset respectively.

**Results of primary and restricted random mating model**

1) Primary random mating model:

Haplotype frequency tests

We found the frequencies of haplotypes B and C in both consort males and fathers were significantly lower than those in the simulated results. We found the frequencies of haplotypes G and H in both consort males and fathers were significantly higher than those in the simulated results. The frequency of haplotype E in consort males tended to be higher but was not significant after Bonferroni correction. The frequencies of haplotypes F and H in consort females tended to be lower and higher respectively, but were not significant after Bonferroni correction. In addition, the frequencies of haplotypes A and G in mothers tended to be lower and higher respectively, but were not significant after Bonferroni correction.

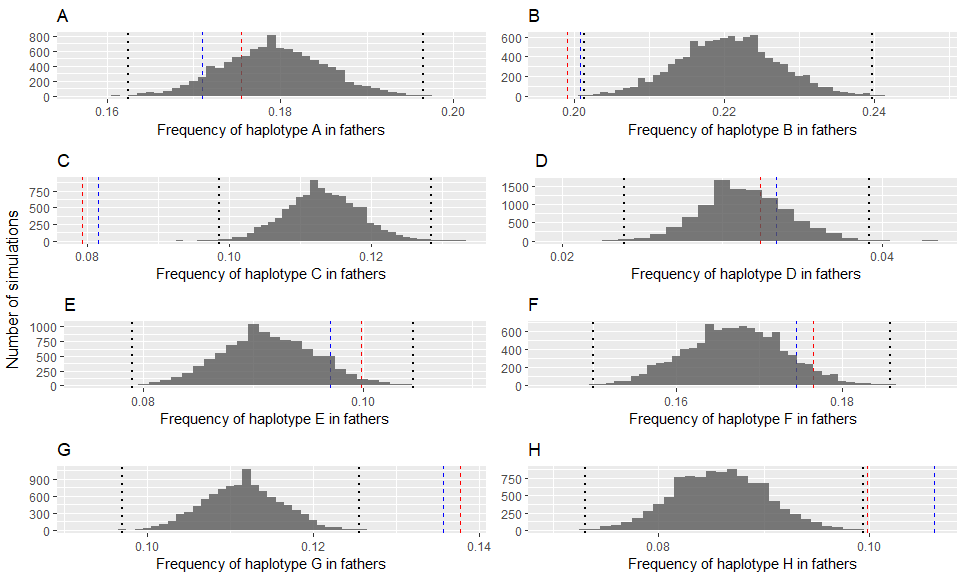

Supplementary figure 5. Results of MHC haplotype frequency tests in fathers under the primary random mating model. Histograms represent the result of simulations with dotted lines representing critical p-value of the distributions after Bonferroni correction. The red and blue dashed blue lines show MHC haplotype frequency in the consort and observed parentage datasets respectively.

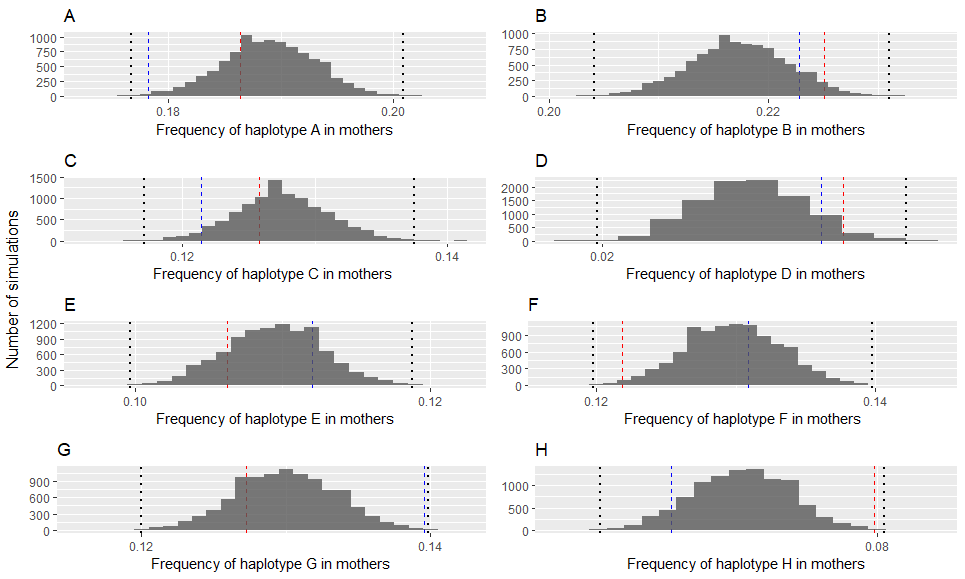

Supplementary figure 6. Results of MHC haplotype frequency in mothers under primary random mating model. Histograms represent the result of simulations with dotted lines represent critical p-value of the distributions after Bonferroni correction. The red and blue dashed blue lines show MHC haplotype frequency in the consort and observed parentage dataset respectively.

Diplotype-based tests

Regarding individual MHC heterozygosity, we found the ratio of homozygote to heterozygote in consort females was significantly higher than expected under the null model. However, the ratio of homozygote to heterozygote in mothers was in line with expectation under the null model. We found the average number of shared haplotypes between parents was significantly lower compared with the null expectation. In addition, the proportion of parents sharing 0 haplotype was significantly higher than the simulated results while the proportion of parents sharing 1 haplotype was significantly lower than the simulated results. However, MHC divergence measured as amino acid sequence differences between parents was in line with expectation under random mating.

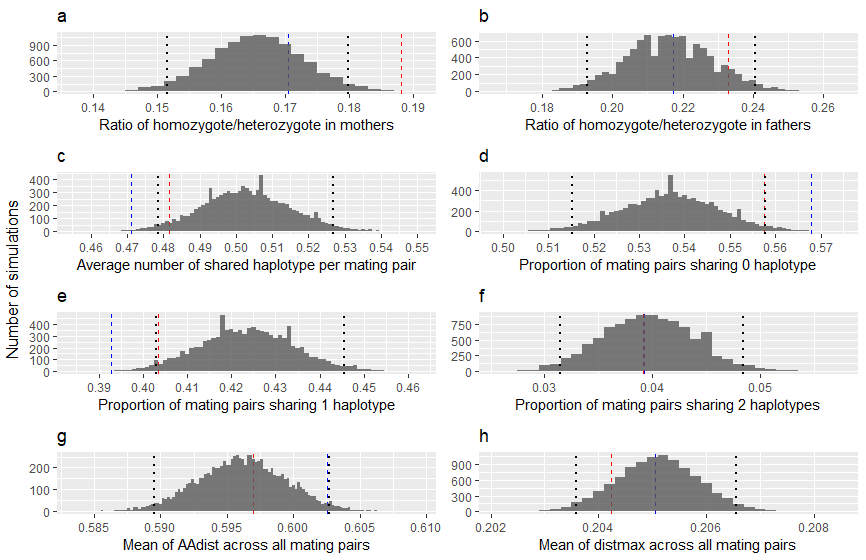

Supplementary figure 7. Results of diplotype tests under under primary random mating model. Histograms represent the result of simulations with dotted lines represent 2.5% and 97.5% tails of the distributions. The red and blue dashed blue lines show indices in the consort and observed parentage dataset respectively.

Genome-wide relatedness tests

We found that the mean and median genomic relatedness between consort pairs and between parents were significantly lower than expected under the null model.

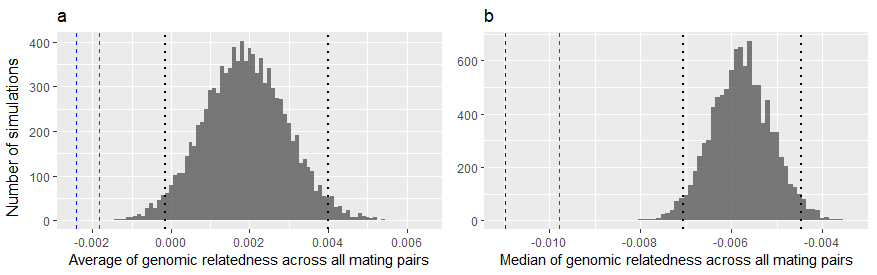

Supplementary figure 8. Results of genome-wide diversity tests under primary random mating model. Histograms represent the result of simulations with dotted lines represent 2.5% and 97.5% tails of the distributions. The red and blue dashed blue lines show MHC divergence in the consort and observed parentage dataset respectively.

2) Restricted random mating model:

As the mating pool only included individuals which actually had offspring in a certain year, MHC haplotype frequency and individual MHC heterozygosity in the simulated results were exactly the same with those in observed parentage dataset. Therefore, only MHC compatibility and genomic relatedness were examined.

We found indices regarding MHC compatibility were in line with expectations under the null model except for average MHC divergence (AAdist) between parents which was significantly higher than that in the simulated results. However, mean and median genomic relatedness between parents were significantly lower compared with simulated results. In addition, mean genomic relatedness between mated pairs were significantly lower compared with simulated results.

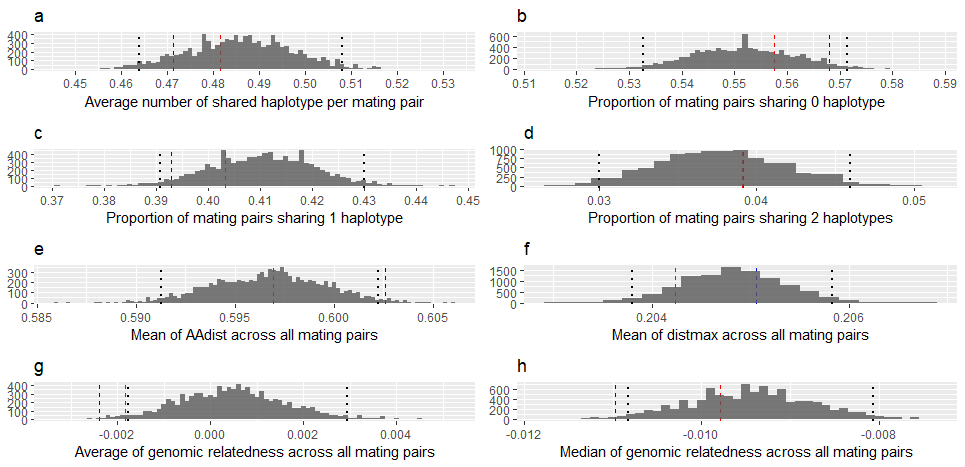

Supplementary figure 9. Results under restricted random mating model. Histograms represent the result of simulations with dotted lines represent 2.5% and 97.5% tails of the distributions. The red and blue dashed blue lines show indices in the consort and observed parentage dataset respectively.
